# “Imaging Translation in Early Embryo Development”

**DOI:** 10.1101/2024.12.09.626398

**Authors:** Pierre Bensidoun, Morgane Verbrugghe, Mounia Lagha

**Affiliations:** Institut de Génétique Moléculaire de Montpellier, University of Montpellier, CNRS-UMR 5535, 1919 Route de Mende, Montpellier 34293 Cedex 5, France; Department of Biochemistry, McGill University, Montréal QC H3G 1Y6, Canada

**Keywords:** Live imaging, SunTag/scFv, translation, smFISH, *Drosophila*, Embryo, developmental biology, cell biology, gene expression

## Abstract

The ultimate output of gene expression is to ensure that proteins are synthesized at the right levels, locations, and timings. Recently different imaging-based methods have been developed to visualize the translation of single mRNA molecules. These methods rely on signal amplification with the introduction of an array of a short peptide sequence (a tag such as SunTag), recognized by a genetically encodable single-chain antibody (a detector such as scFv). In this chapter, we discuss such methods to image and quantify translation dynamics in the early *Drosophila* embryo and provide examples based on a *twist-32XSunTag* reporter. We outline a step-by-step protocol to light-up translation in living embryos. We also detail a combinatorial strategy in fixed samples (smFISH-IF), allowing to distinguish single mRNA molecules engaged in translation.

## 1 Introduction

During embryogenesis, gene expression is tightly controlled in space and time to allow the adoption of specific and reproducible cellular identities. A major functional product governing these decisions is proteins. Thus, to obtain a comprehensive view of how gene expression is controlled in space and time, it is important to monitor the flow of information along the central dogma of molecular biology. While imaging methods have been developed more than 20 years ago to image mRNA synthesis, an equivalent method was only recently deployed to visualize the translation of single molecules of mRNA in live cells (1–8). Different fluorescent microscopy strategies allow to image translating mRNAs at the single molecule level in living cells. Most approaches revolve around a similar signal amplification system whereby an mRNA of interest bears sequences encoding for repetitions of a short peptide, recognized by a genetically encoded fluorescent detector -usually a soluble antibody-co-expressed in cells. The detectors bind to the nascent peptides emerging from the ribosome exit sites, allowing to pinpoint translation events and to grasp their dynamics. In this chapter, we discuss imaging-based signal amplification techniques to monitor translation dynamics in *Drosophila* embryos. We particularly focus on the SunTag approach where nascent peptides derived from the yeast Gcn4 protein^1^ are recognized by a recombinant single-chain variable fragment (scFv) antibody fused to a fluorescent protein (FP) (see note 1). To amplify the signal provided by translation events, mRNAs of interest contain typically 32 or 24 Suntag tandem repeats (6-8 – see note 2).

The Suntag array is usually inserted in frame, within the 5’ untranslated region (5’UTR) of the mRNA of interest which facilitates the detection of early translation events. To simultaneously visualize individual mRNAs together with nascent peptides in living cells, the SunTag/scFv system is generally combined with the MS2/MCP system (8-11 –see note 3-4). In this method, transcripts can be labeled with MS2 repeats and detected with an MCP detector protein fused to a fluorescent protein. To avoid perturbing translation initiation or ribosome scanning, MS2 tags are generally inserted in the 3’UTR region (12).

Alternatively, translation events can be detected in fixed samples by simultaneously labeling single mRNA molecules with single-molecule fluorescent in situ hybridization (smFISH) using SunTag probes in combination with protein labeling by immunofluorescence (using an anti- GCN4 antibody recognizing the SunTag peptides (13 – see note 6-7).

These methods were successfully implemented by our group and others to characterize the translation modalities and kinetics of critical developmental mRNAs such as the zygotically expressed *hunchback* (7), *twist* or the maternal *nanos* mRNAs (8). In this chapter, we describe a detailed protocol to prepare and mount fly embryos for imaging *twist* mRNA translation both in fixed (Figure 3) or living embryos (6,7,13 – Figure 1, Figure 2).

**Figure 1:**
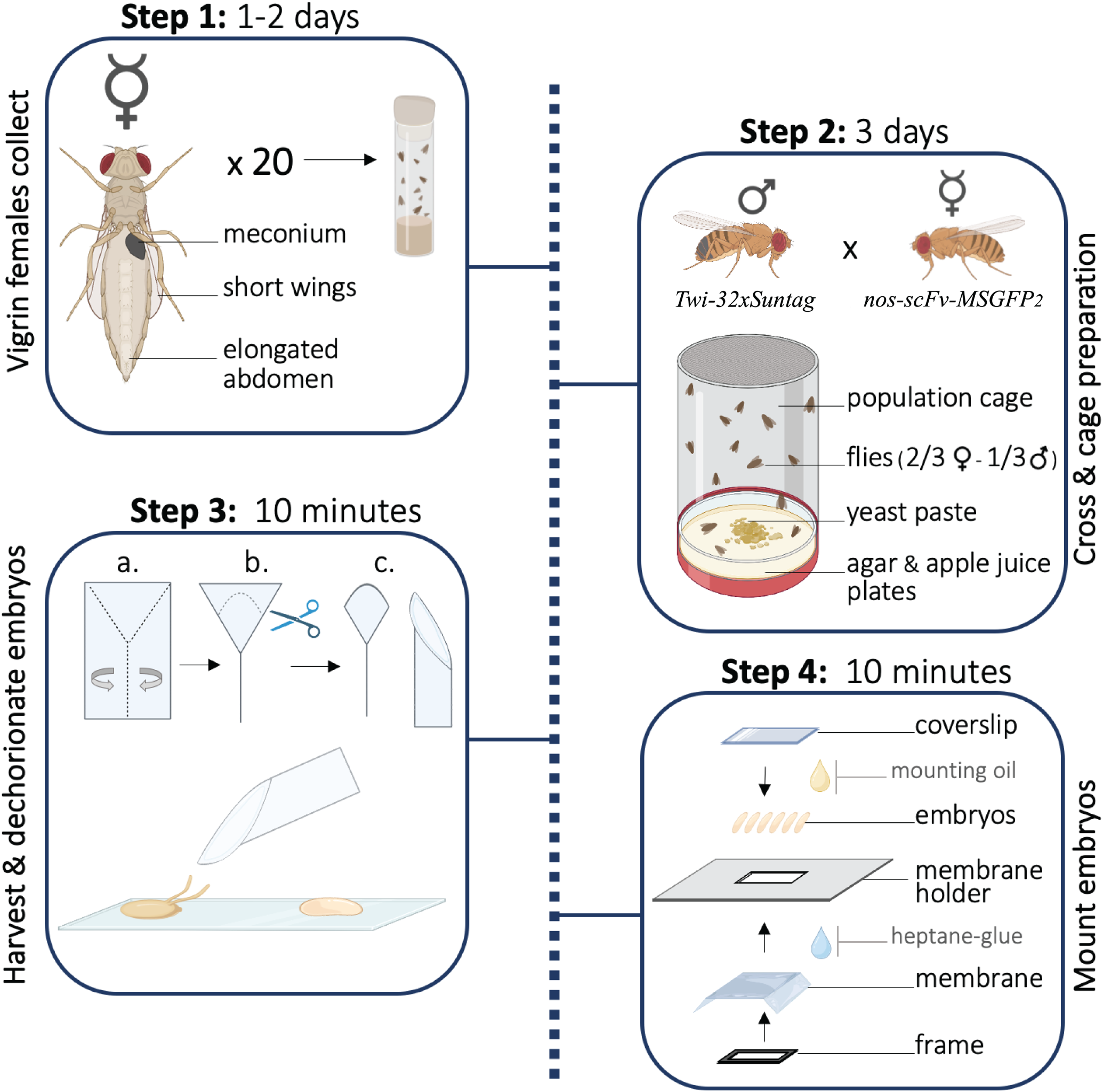
Overview of collecting and mounting embryos for live imaging of translation. Schematic indicates the timing, quantity of material and genotype of the fly stocks are indicated.

**Figure 2:**
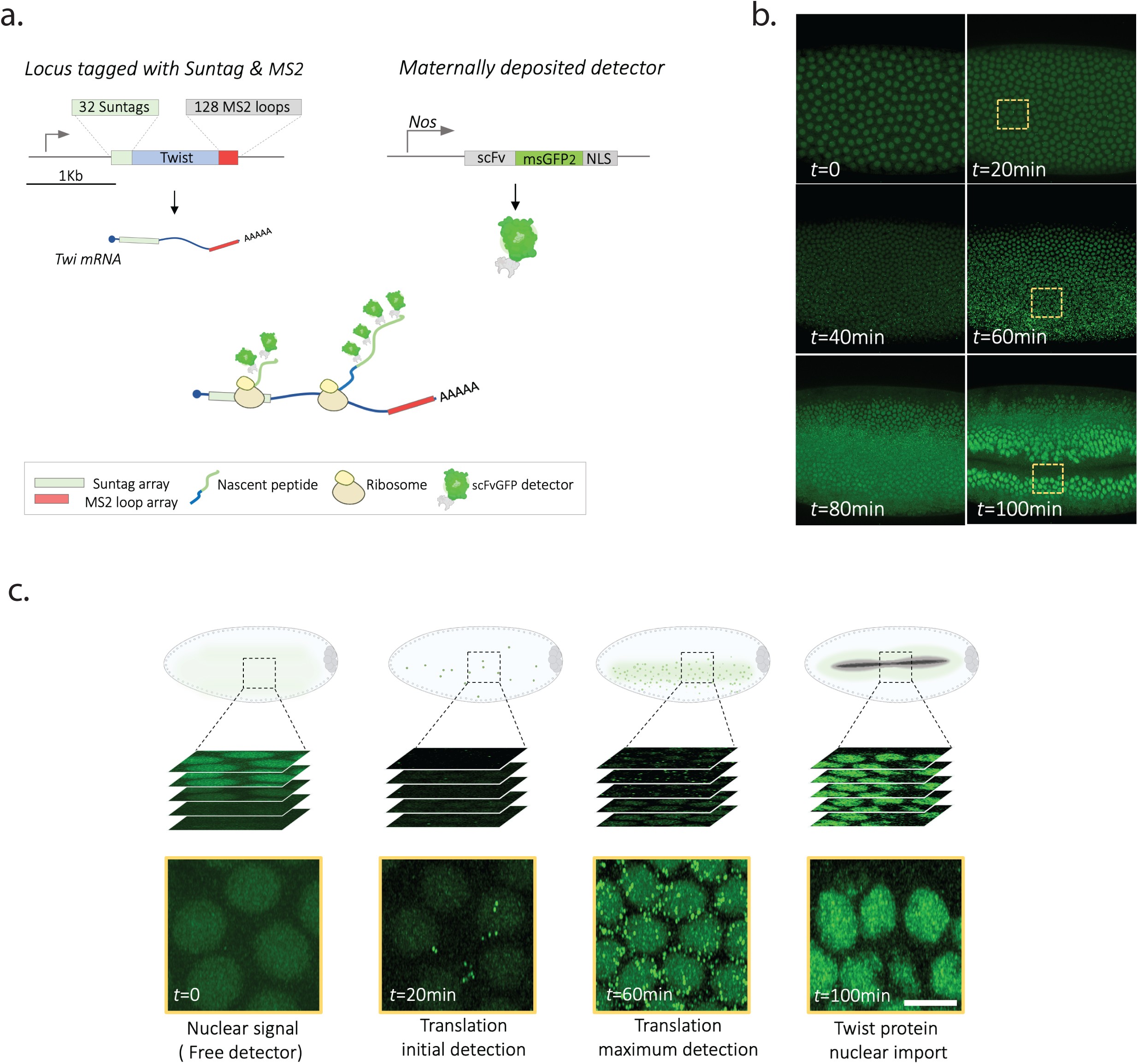
Imaging twist translation in living embryos. a. Principle of the SunTag/scFv labeling method, the example of *twist-SunTag-MS2* allele. b. Still images from a movie obtained by confocal imaging of a living *twist-Suntag; scFv-NLS-MSGFP2* from early nc14 to gastrulation. c. Zoomed images of the insets indicated as yellow rectangles in b depicting typical signals obtained at various timings. Imaging *twist-SunTag* with an scFv detector typically shows 3 types of signals: free detector labeling (nuclear); cytoplasmic dots and mature protein imported in nuclei (Twist is a transcription factor (Scale bar=6µm).

**Figure 3:**
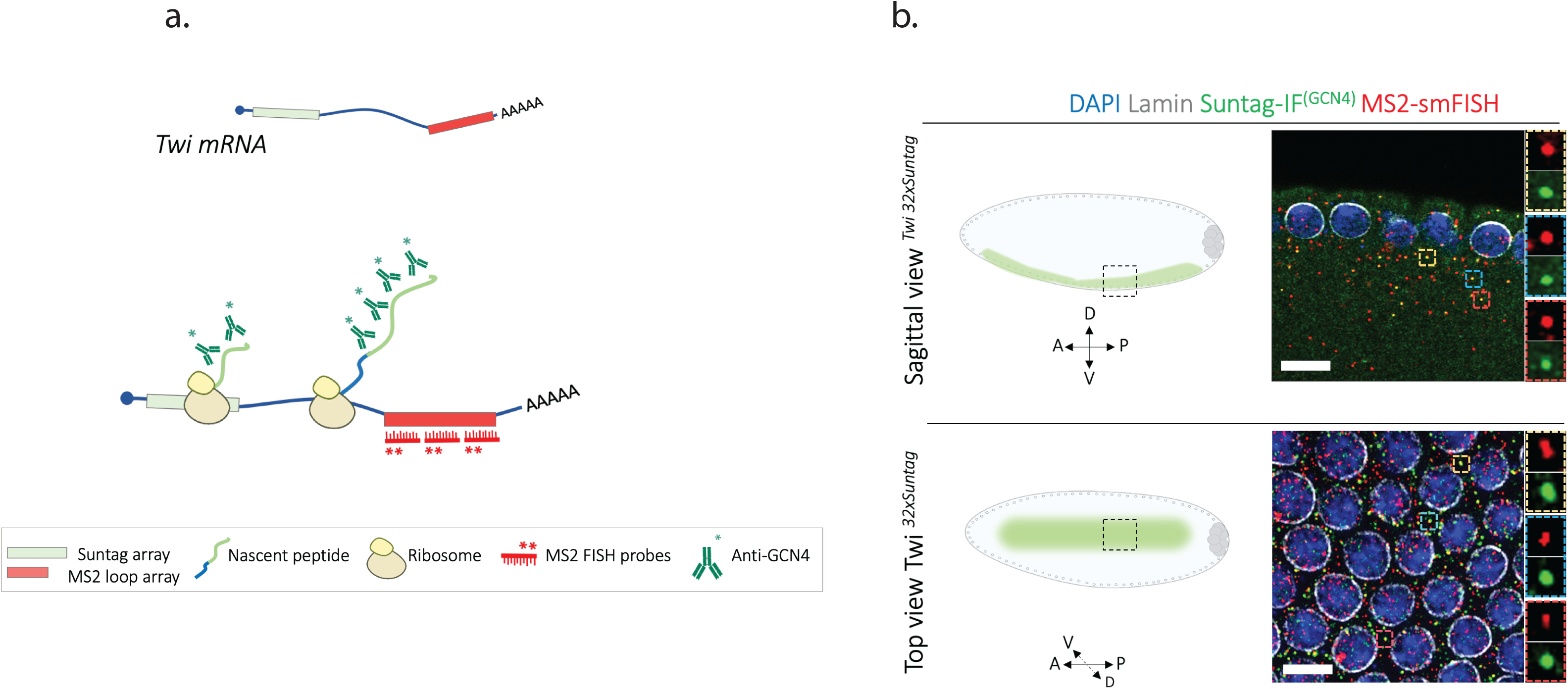
Imaging translation of single molecules of mRNAs in fixed embryos. a. Schematic of the rationale to directly detect translation from a SunTag reporter (in the absence of detector proteins). b. Confocal images of smFISH-IF experiment from two orientations: sagittal (top panel) and top view (bottom panel). MS2 probes signal is shown in red, staining of lamins, GCN4, and Dapi are shown in grey, green, and blue respectively. Zoomed examples of colocalizing MS2 and GCN4 signals are shown on the cropped right panels (Scale bar=6µm).

## 2 Materials

In this method, we describe sample preparation and imaging set-up to examine the translation of *twist* mRNAs in pre-gastrulation embryos throughout the timing of zygotic genome activation, from the nuclear cycles (nc) 12 to nc 14 (see note 1-4). The endogenous *twist* allele was modified with 32x SunTag repeats and 128 MS2 stem loops by CRISPR- cas9 technology. For a typical live imaging experiment, the modified *twist* allele is provided by males, while the fluorescent detector proteins, termed scFv-msGFP2, are maternally transmitted (Figure 1). This maternal pool of detectors ensures that enough scFv-msGFP2 is present in the embryo during the earliest steps of development (see note 1).

### 2.1 Fly lines

A typical experiment of live imaging requires two parental fly stocks:

1. The SunTag detector scFv is expressed from a transgenic line, *scFv- msGFP2*, controlled by the *Nanos* (*nos*) promoter and proximal enhancer, inserted in chromosome III by PhiC targeted insertion (13). Other detector lines can be used and are listed in Table 1.
2. The ‘Tag’ stock, expressing the SunTag and MS2 arrays: *twi_SunTag_MS2_CRISPR*.

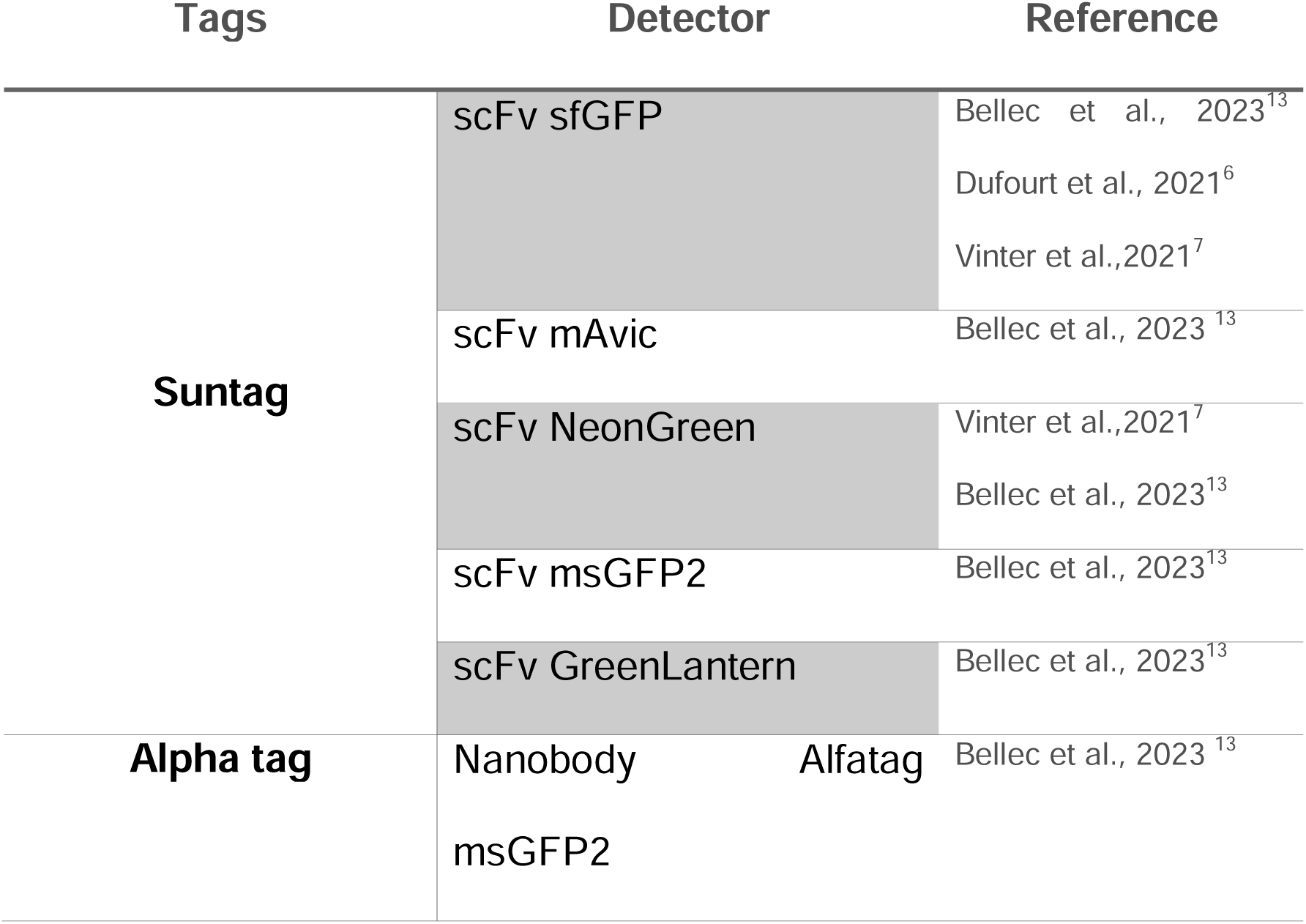

### 2.2 Tags and detectors: Sun tag, Alpha tag

To unravel translation dynamics and translation heterogeneities between mRNA species, several versions of the nascent peptide labeling system have been successfully deployed in the fly embryo. Analogously to the SunTag approach, the Alpha-tag strategy uses two components: multiple Alpha tag peptides recognized by an existing free binder, the anti-Alpha nanobody (13–14). The pairs of tag-detectors successfully used in our laboratory and others (6,7,13) for early translation imaging in fly embryos are listed in Table 1.

1. To detect translation events in fixed samples by immunofluorescence we use a commercial version of a mouse GCN4 antibody (Novus Biologicals ; C11L34)
2. GFP-conjugated anti-mouse secondary antibody (ThermoFisher; A32766TR) (see note 6).
3. Our FISH probes set hybridizing on the MS2 repeats are 19-23 mer antisense DNA oligos designed with Stellaris^TM^ Probe designer algorithm. We recommend using at least 20 probes per transcript which can be ordered already conjugated with dyes such as Cy3, Cy3.5, or Cy5 from any provider (see note 7).
4. To estimate the developmental stage and orient each embryo, we rely on the DAPI staining (Invitrogen; 1825107) and/or nuclear envelope labeling with a lamin antibody (ADL67.10).

### 2.4 Media, Solution, Consumables and Buffers Fly breeding

1. Apple juice 5% agar plates (35mn)
2. Small population cages (35mn)
3. Active dry yeast
4. Paintbrush (≤2.4mm)

For live imaging

1. Transparent double-sided tape
2. Breathable Biofoil film.
3. Heptane glue.
4. 10X Oil.
5. 1.5µm coverslips.
6. Glass slides.
7. Membrane holder/ Plastic embryo holder.
8. Membrane holder frame
9. Sieves or mesh basket (with nylon mesh pores of 100µm)

Specific for fixed samples

1. Bleach
2. Formaldehyde
3. Methanol
4. Ethanol
5. Tween20
6. Rnase inhibitor
7. Phosphate buffer saline (PBS)
8. BSA
9. Formamide
10. 20x SSC
11. Dextran sulfate
12. VRC
13. MilliQ water
14. tRNA
15. prolong gold media

#### Buffers

PBST-1X (1L) : Sodium Chloride (NaCl) 137 mM , Potassium Chloride (KCl) 2.7 mM, Disodium Phosphate (Na_2_HPO_4_) 10 mM, Monopotassium phosphate (KH_2_PO_4_) 1.8 mM, Tween 20 0.1%, MilliQ water to reach 1 L.

*The PBST can be stored at room temperature

*Add the RNasin and the BSA fresh before use. For 10 mL of PBST, we add 10µL of RNAsin (2500U/µl) and 1 mL of BSA (50mg/mL).

Wash buffer (10 mL) : SSC 2x , Formamide 10%, MilliQ water to reach 1 L.

*Wash buffers are always made fresh.

Hybridization buffer : Formamide 10%, SSC 2x , tRNA 0.36mg/ml, Dextran sulfate 5.5%, Stellaris probe 50nM, VRC (Vanadyl-ribonucleoside complex) 2nM, BSA 0.9mg/ml, Primary antibody anti GCN4 0.5µg/ml, MilliQ water to reach 300 µL.

*New hybridization buffer is always prepared fresh and protected from light. However, the hybridization buffer can be reused for new FISH experiments at least two more times. To do so, the buffer is carefully removed from the tube (avoiding aspirating embryos) and kept at -20°C.

### 2.5 Equipment

1. Micropipettes.
2. Standard shakers and table centrifuges.
3. 37°C dry incubator
4. Rotating wheel
5. Binocular magnifier with a non-heating light source

## 3 Methods

### 3.1 Live imaging

#### 3.1.1 Fly cross

1. Collect ∼20 virgin females carrying the fluorescent-free detector scFv- msGFP2 from amplification bottles. This step usually takes 1-2 days.
2. Rehydrate the active dry yeast to obtain a thick yeast paste. Place half a teaspoon of yeast paste on a 35mm apple juice plate to assemble the bottom of the population cage. Flies will lay their eggs on the surface of the apple juice plate.
3. Females are placed in a population cage, with ∼10 males carrying the SunTag/MS2 tagged *twist* gene. Cages are kept in an incubator with day-night cycles at 25°C.
4. The apple juice plate with fresh yeast paste is changed every day for at least 3 days before we image.

#### 3.1.2 Collecting and mounting embryo

1. Prepare a mounting device with 4cm^2^ of a breathable Biofoil film maintained into the membrane holder and the frame as described in Figure 1 (Step 4). The breathable Biofoil hydrophobic side should be facing up. Of note, the hydrophobic side of the Biofoil can be identified easily as permanent maker ink does not properly write on it.
2. With a Pasteur pipette, drop 100µl of heptane glue on the hydrophobic side and let it dry for a few minutes.
3. To ensure that you harvest pre-gastrulating embryos, change the apple juice plate with fresh yeast in the morning. To enrich for early embryos ( < nc 12 - nc 13), harvest 3h after switching the plate. Conversely, to enrich for nc14 embryos and gastrulating embryos we typically let the females lay for 2h, then remove the collection plate which is kept on the bench for 2 hours.
4. On a microscope slide, add double-sided tape and gently deposit ∼20 embryos with a clean paintbrush.
5. Fold a piece of double-sided tape into a spoon shape as described in Figure 1 (Step3 a;b). Adjust the end of the folded tape to obtain a rounded form (Step 3 c).
6. Using the double-tape spoon, roll the embryos on the slide to remove the chorion. Dechorionated embryos are glossy and transparent. To avoid desiccating the embryos, this step should be completed as fast as possible.
7. Transfer each embryo onto the Biofoil membrane on the mounting slide as described in Figure 1 (Step 4). At this step, you can adjust the position of the embryos to facilitate the imaging by rolling them with the double tape spoon: the ventral region is slightly concave.
8. Add a droplet (100µl) of 10X oil onto the coverslip and place it over the embryos. Carefully press the four corners of the coverslip.
9. Clean the oil swelling from the sides of the coverslip with a towel and 70% ethanol. At this point, the coverslip should be fairly stable and the embryos are ready to image.

#### 3.1.3 Data acquisition for live imaging

1. To image live embryos, use an inverted confocal microscope LSM 980 from Zeiss with an Airyscan 2 module. This microscope allows monitoring the translation with a relatively low amount of light to limit phototoxicity and photobleaching. Live imaging of translation is also possible with other types of confocal microscopes or with light sheet microscopy such as MUVISPIM^6^) (see note 5).
2. We generally use a 40 X plan apo 1.3NA objective with oil immersion. Therefore, a droplet of oil ∼40µl is deposited onto the objective before imaging.
3. For consistent measurements between acquisitions, control the laser power using a laser power meter before each imaging session. The typical laser power employed lies in the range of 2% of the maximum laser power (∼6 µW). If the maximum laser power decreases, compensate by increasing the percentage of laser power utilized to illuminate the sample.
4. The Airyscan module needs to be calibrated before each acquisition. This can be achieved with a channel using the continuous mode in Airyscan. Calibration is accurate in a channel providing enough signal to measure a range of 60 levels of grey in detector view. However, this calibration procedure does not necessarily need to be performed in the channel of interest.
5. In the absence of a nuclear or a membrane marker, it can be challenging to locate the embryos. However, once nuclei migrate to the surface, the fluorescence emanating from the free scFv-msGFP2 detector pool is sufficient to identify blastoderm embryos as the detector contains a nuclear localization signal (Figure 2b – see note 1).
6. The settings applied to image *twist* translation are listed in the metadata table 2. To be able to identify as many translation events as possible in 3 dimensions, acquire 20 stacks per frame (see note 4 and 5).
7. Depending on the biological question, acquisition can last from 1 to 3 hours. Live imaging is generally manually stopped once the ventral furrow is formed. *twist* mRNA encodes for a transcription factor. Therefore, mature Twist proteins labeled with SunTag are imported into nuclei, and Twist protein steady states appear in the last part of the acquisition as bright fluorescent nuclei within the mesoderm. Thus, a typical movie of *twist* translation will show three types of signals (Figure 2c): free scFv-msGFP2 (labeling nuclei); bright cytoplasmic dots corresponding to translation foci, and fluorescent nuclei within the mesoderm (mature Twist protein after nuclear import; see note 1).

**Metadata Table 2.**
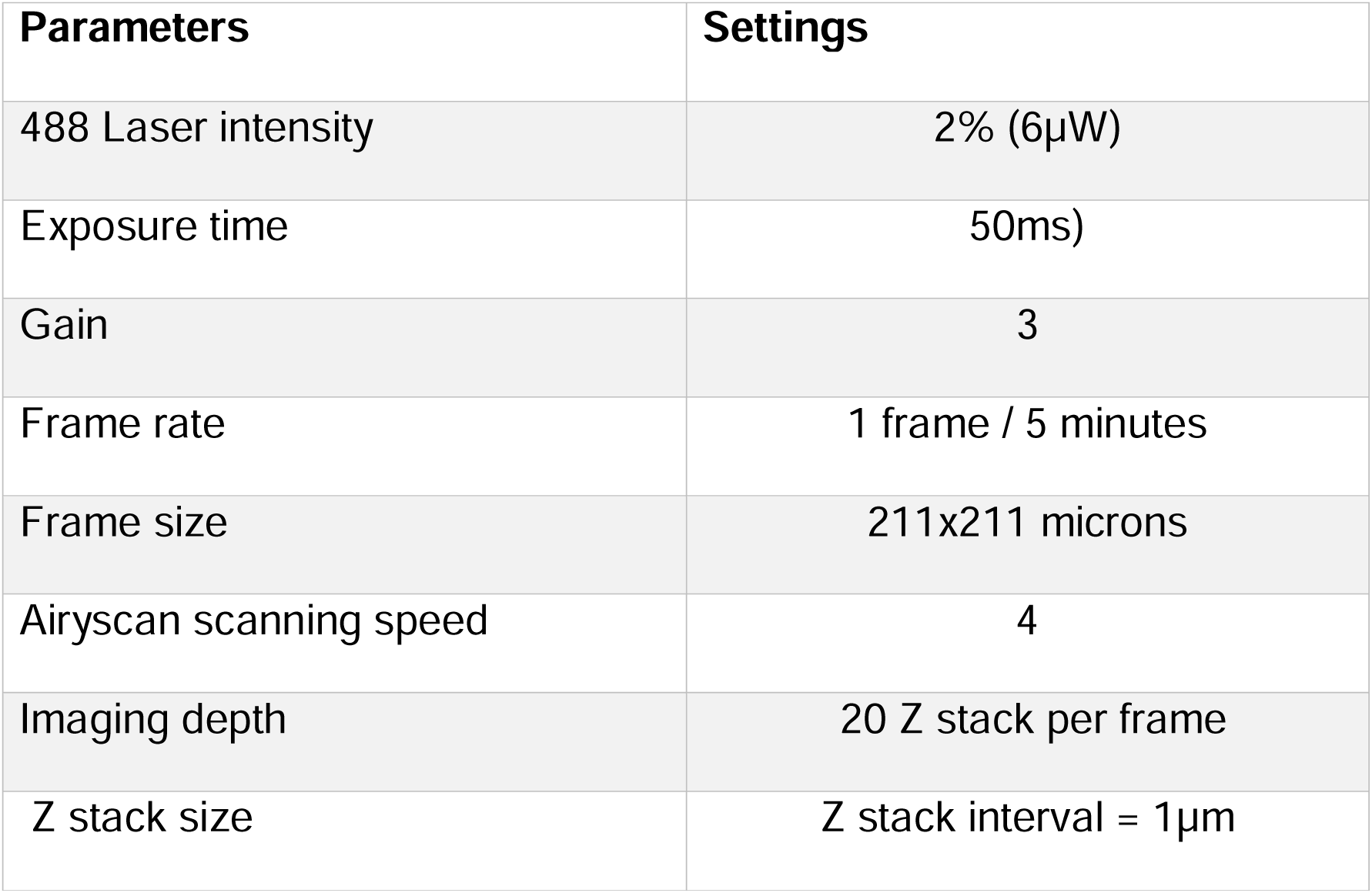

### 3.2 Fixed samples

#### 3.2.1 Fly cross and embryo harvest

1. The cross is set up as described in section 3.1.1.
2. To make sure that enough embryos are collected for the FISH-IF experiment, do at least 5 rounds of 0-5h embryo harvests. Each time the plate is cleaned with distilled water to remove the remaining embryo and yeast paste.
3. Plates can be recycled between the harvesting rounds for embryos generated from the same cross.

#### 3.2.2 Embryo fixation

1. After a brief rinse with clear water, assemble the basket with the mesh.
2. Gently pour water on the apple juice plate and carefully remove the embryos from the plate with a paintbrush to transfer them into the basket.
3. To chemically remove the chorion from the embryos, immerse the net of the basket in bleach for 3min.
4. Wash thoroughly the embryos with water
5. With a paintbrush, place the embryos in a 2ml Eppendorf tube containing 1ml of 5% Formaldehyde, and 50% Heptane.
6. Agitate the embryos for 25min using a horizontal shaker at 400rpm.
7. Remove the bottom phase (Formaldehyde layer) and add 500ul of 100% methanol.
8. Vigorously shake the tube by hand for 1min.
9. Remove the top layer and the interphase.
10. Rinse briefly 3 times with 100% methanol.
11. At this step, embryos can be stored at -20°C.

#### 3.2.3 Single-molecule FISH and IF

1. Wash embryos in 1ml 50% ethanol and 50% methanol and place them for 5min on a rotating wheel.
2. Rinse briefly twice with 1mL of 100% ethanol.
3. Wash for 5 min with 1mL of 100% ethanol on a rotating wheel. Repeat this step twice.
4. Rinse briefly in 100% methanol.
5. Wash 5min with 1mL of 100% methanol on a rotating wheel.
6. Rinse briefly twice in 1mL of PBST-RNAsin buffer.
7. Wash the embryos 4 times for 15min in 1mL PBST with RNAsin + 0.5% BSA.
8. Wash 20min in 1mL of wash buffer.
9. Prepare the hybridization buffer. As conjugated probes are light-sensitive, keep the hybridization buffer in the dark.
10. Remove the wash buffer from the embryos and add the hybridization buffer (see note 6).
11. Incubate the embryos overnight in the hybridization buffer at 37°C.
12. Wash twice the embryos in wash buffer pre-warmed at 37°C.
13. Wash the embryo in wash buffer with 0.5% BSA and 1.5 µL of secondary antibody (anti-mouse 555 1/500^e^) for 45 min at 37°C.
14. Add 0.3 µL of DAPI (1/1000^e^) and leave at 37°C for 15min.
15. Wash three times in 2X SSC 0.1% tween. After the last wash, take out the buffer and remove as much solution as possible.
16. Cut the extremity of a 200 µL tip and add 110 µl of mounting prolong diamond antifade media to the embryos.
17. Disperse the embryos over the slide, carefully using a low bind tip.
18. Once the embryos are well spread out, add the coverslip and let the slide dry for at least 24h in a box protected from light.
19. The slide can be directly imaged or kept at 4°C.

#### 3.2.4 Data acquisition

1. To image SmFISH-IF experiments, we employed an inverted confocal microscope LSM 880 from Zeiss with a fast Airyscan module. This microscope is equipped with a 405 diode together with Argon laser 458, 488, and 514.
2. Use a 40X N.A 1,4 plan apochromat oil objective immersed in ∼40µl of oil before each acquisition.
3. The laser power is checked and adjusted as described in the 3.1.3 live imaging section, step 3.
4. The Airyscan is calibrated as described previously in the 3.1.3 live imaging section, step 4.
5. Embryos are selected depending on the stages using the DAPI signal.
6. To keep a trace of the embryo orientation,take a snap image of the embryo with a 0.6 Zoom.
7. A smaller area using a 4.0 zoom is then acquired using the parameter described in metadata table 3.
8. Depending on the orientation the localization of the mRNA and its translation can be imaged from the top or in the sagittal plane as shown in Figure 3b. Both angles can be used to examine mRNA localization and topological heterogeneities of translation.

**Metadata Table 3.**

| Parameters | Settings |
| --- | --- |
| 405 Laser intensity | 2% (1μW) |
| 488 Laser intensity | 0.8% (3μW) |
| 566 Laser intensity | 1% (4μW) |
| 647 Laser intensity | 2% (6μW) |
| Exposure time | (50ms) |
| Gain | 3 |
| Frame size | 211x211 microns |
| Airyscan scanning speed | 4 |
| Imaging depth | 20 Z stack per frame |
| Z stack size | Z stack interval = 1μm |

## 4 Notes

1. The signal amplification system described in this method relies on the presence of a pool of free detector proteins, fused to a fluorescent protein. Determining the right free detector scFv- msGFP2 concentration is key to the success of translation imaging. On the one hand, this pool needs to be as low as possible in the cytoplasm to ensure a good signal-to-noise ratio when translation foci appear, but on the other hand, levels of detectors should be saturating to avoid their titration when high amounts of transcripts are translated. To maintain low levels of free detectors in the cytoplasm, you can include a nuclear localization signal to both Suntag and Alpha detectors. To test whether detector levels are not limiting, systematically perform FISH-IF with an antibody against the tag-peptide (GCN4 for SunTag and anti-Alpha for ALFA-Tag). Comparing the percentages of mRNA engaged in translation in the absence of a detector (direct IF, Figure 3) with those obtained in the presence of a detector (in fixed cells) informs on the level of potential titration. If titration is an issue, one can increase the levels of detector proteins, with strong drivers (UAS-scFv- MSGFP2 – 13). However, increasing detector levels may significantly enhance the background caused by the unbound fraction of detectors as well as promote aggregate formation. Alternatively, one may consider reducing the number of transcripts bearing the tag (SunTag or ALFA tag repeats), using for example expression from transgenes instead of Crispr (for example *twist*-Enhancer-SunTag transgene – 6).
2. In this method, we describe the strategy to follow *twist* mRNA translation using 32 SunTag repeats. Translation can be monitored with fewer SunTag repeats as 24 SunTags repeats were sufficient to reveal the translation of *hunchback* mRNA (7).
3. Adding exogenous repetitive motifs such as MS2 or SunTag sequences may interfere with mRNA turnover. The relative abundance can be used as a proxy for mRNA half-life and tagged *versus* untagged mRNA levels can be estimated and controlled by mRNA FISH, Northern blot, or RT-qPCR. Over- stabilization of transcripts is sometimes observed when detectors are also co-expressed in cells and may correlate with the formation of aggregates containing both mRNAs and detectors. (9,15). In this chapter, we use a recent version of the scFv-msGFP2 detector, for which we never observed aggregation (13).
4. Quantitative imaging requires isolating single mRNA molecules or individual translation sites. For strongly expressed mRNAs, or highly localized transcripts, the domain occupied by the transcripts appears crowded, and pinpointing single molecules is challenging. To circumvent this limitation, you can apply similar strategies (described in note 1) using weaker promoters to drive the expression of our gene of interest or work with embryos where only one copy of the allele contains the tags.
5. Live imaging of translation spots can be analyzed to extract translation kinetics such as particle movements, velocity, or diffusion coefficient (16–17). Moreover, using similar approaches and models to those developed to quantify transcription dynamics, temporal correlation analyses can be employed to infer kinetic parameters of translation (elongation speed, initiation rate – 6,18-21). However, these approaches require ultra-fast imaging (typical temporal resolution on the order of 500ms/frame). Indeed, extracting kinetics requires the observation of particles for long periods of time (typical 300ms for *twist*-SunTag – 6), which is difficult to obtain due to the limitations of bleaching and 3D diffusion (mainly in Z). For a better temporal resolution, the acquisition speed can be increased by reducing the size of the region of interest or using lattice-light sheet microscopy (22).
6. Some antibodies are not compatible with the FISH protocol and do not provide any good IF staining when they are incubated with fixed embryos in the FISH probes hybridization buffer. Here, once the hybridization of the probes is finished (see section 3.4.3 step 12), embryos are washed 1h in the wash buffer and 15 min in the last wash buffer. Samples are rinsed 3 times in PBT and blocked in PBT 5% BSA for 40min. The antibody is added directly to the PBT 5% BSA at the working concentration and samples are kept in the dark overnight. Samples are washed in PBT 5% BSA 3 times for 5min before the incubation with secondary antibody for 40 min in PBT 5% BSA. Finally, samples are rinsed 3 times with PBT and we proceed as described in section 3.2.3, step 14.
7. The MS2-MCP system has been optimized to tag and track individual mRNAs in fly embryos. In the protocol presented in this chapter, the MS2 arrays serve as a label for FISH probes hybridization. This highly repetitive region is recognized by a large number of FISH probes (>20 DNA oligos) and provides good smFISH signals. Alternatively, one can also use probes against the Suntag or 16xAlpha tag arrays with similar efficiency as the MS2 FISH. Using repeated arrays of exogenous sequences as beacons for probes hybridization allows for standardizing the detection efficiency between different MS2, Suntag, or Alpha-tagged mRNA species (Figure 3).

## Acknowledgment

We thank members of the Lagha team for insightful discussions and particularly A.Palandri for her technical support in fly handling and advice on *Drosophila* genetics and for troubleshooting specific steps of the methods. Initial work on translation was supported by an HFSP CDA grant to ML and the ANR MemoRNP. PB and MV are supported by the ERC LightRNA2Prot grant. ML is sponsored by the CNRS.

## Notes

### Competing Interest Statement

The authors have declared no competing interest.

